# Developmental transcriptomes of the sea star, *Patiria miniata*, illuminate the relationship between conservation of gene expression and morphological conservation

**DOI:** 10.1101/573741

**Authors:** Tsvia Gildor, Gregory Cary, Maya Lalzar, Veronica Hinman, Smadar Ben-Tabou de-Leon

**Affiliations:** Department of Marine Biology, Leon H. Charney School of Marine Sciences, University of Haifa, Haifa 31905, Israel.; Bionformatics Core Unit, University of Haifa, Haifa 31905, Israel.; Departments of Biological Sciences and Computational Biology, Carnegie Mellon University Pittsburgh, PA 15213, USA

## Abstract

Evolutionary changes in developmental gene expression lead to alteration in the embryonic body plan and biodiversity. A promising approach for linking changes in developmental gene expression to altered morphogenesis is the comparison of developmental transcriptomes of closely related and further diverged species within the same phylum. Here we generated quantitative transcriptomes of the sea star, *Patiria miniata* (*P. miniata*) of the echinoderm phylum, at eight embryonic stages. We then compared developmental gene expression between *P. miniata* and the sea urchin, *Paracentrotus lividus* (~500 million year divergence) and between *Paracentrotus lividus* and the sea urchin, *Strongylocentrotus purpuratus* (~40 million year divergence). We discovered that the interspecies correlations of gene expression level between morphologically equivalent stages decreases with increasing evolutionary distance, and becomes more similar to the correlations between morphologically distinct stages. This trend is observed for different sub-sets of genes, from various functional classes and embryonic territories, but is least severe for developmental genes sub-sets. The interspecies correlation matrices of developmental genes show a consistent peak at the onset of gastrulation, supporting the hourglass model of phylotypic stage conservation. We propose that at large evolutionary distance the conservation of relative expression levels for most sets of genes is more related to the required quantities of transcripts in a cell than to conserved morphogenesis processes. In these distances, the information about morphological similarity is reflected mostly in the interspecies correlations between the expressions of developmental control genes.

**Author summary:** Understanding the relationship between the interspecies conservation of gene expression and morphological similarity is a major challenge in modern evolutionary and developmental biology. The Interspecies correlations of gene expression levels have been used extensively to illuminate these relationships and reveal the developmental stages that show the highest conservation of gene expression, focusing on the diagonal elements of the correlation matrices. Here we generated the developmental transcriptomes of the sea star, *Patiria miniata*, and used them to study the interspecies correlations between closely related and further diverged species within the echinoderm phylum. Our study reveals that the diagonal elements of the correlation matrices contain only partial information. The off-diagonal elements, that compare gene expression between distinct developmental stages, indicate whether the conservation of gene expression is indeed related to similar morphology or instead, to general cellular constraints that linger throughout development. With increasing evolutionary distances the diagonal elements decrease and become similar to the off-diagonal elements, reflecting the shift from morphological to general cellular constraints. Within this trend, the interspecies correlations of developmental control genes maintain their diagonality even at large evolutionary distance, and peak at the onset of gastrulation, supporting the hourglass model of phylotypic stage conservation.

## Introduction

Embryo development is controlled by regulatory programs encoded in the genome that are executed during embryogenesis, which usually last days to months [1]. Genetic changes in these programs that occur in evolutionary time scales - millions to hundreds of million years - lead to alterations in body plans and ultimately, biodiversity [1]. Comparing developmental gene expression between diverse species can illuminate evolutionary conservation and modification in these programs that underlie morphological conservation and divergence. To understand how these programs change with increasing evolutionary distances it is important to study closely related and further divergent species within the same phylum.

The echinoderm phylum provides an excellent system for comparative studies of developmental gene expression dynamics. Echinoderms have two types of feeding larvae: the pluteus-like larvae of sea urchins and brittle stars, and the auricularia-like larvae of sea cucumbers and sea stars [2]. Of these, the sea urchin and the sea star had been extensively studied both for their embryogenesis and their gene regulatory networks [3–11]. Sea urchins and sea stars diverged from their common ancestor about 500 million years ago, yet their endoderm lineage and the gut morphology show high similarity between their embryos [6]. On the other hand, the mesodermal lineage diverged to generate novel cell types in the sea urchin embryo (Fig. 1A, [8, 12]). Specifically, the skeletogenic mesoderm lineage, which generates the larval skeletal rods that underlie the sea urchin pluteus morphology, and the mesodermal pigment cells that give the sea urchin larva its red pigmentation (arrow and arrowheads in Fig. 1A). Thus, there are both morphological similarities and differences between sea urchin and sea star larval body plans that make these classes very interesting for comparative genetic studies.

**Figure 1.**
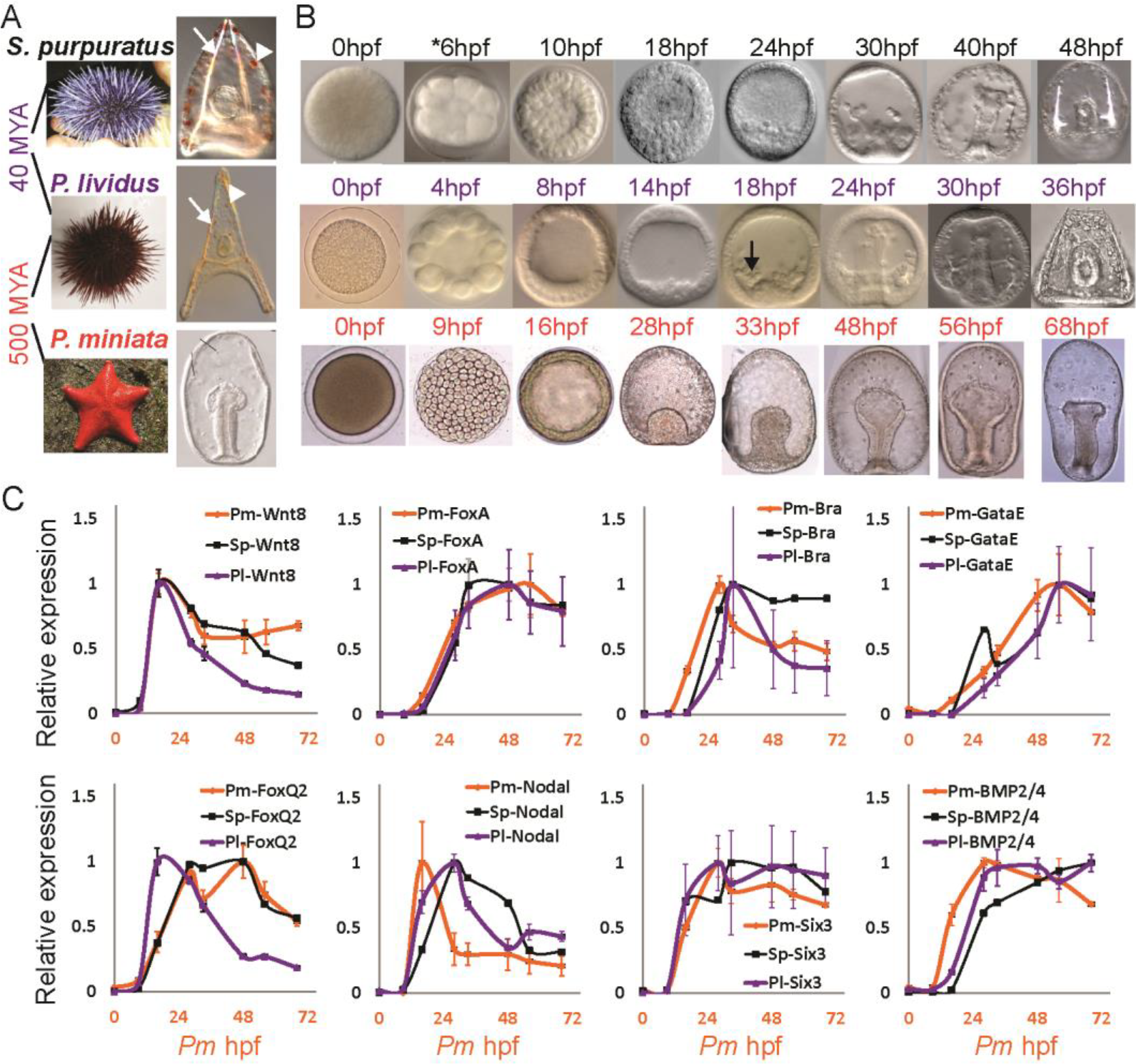
Developmental time points studied and examples for gene expression profiles on the three species. (A) Images of adult and larval stage of *P. miniata*, *P. lividus* and *S. purpuratus* Arrows point to the sea urchin skeletogenic rods and arrow heads point to the sea urchin pigments. (B) Images of *P. miniata*, *P. lividus* and *S. purpuratus* embryos at the developmental stages that were studied in this work. Time point 6hpf in *S. purpuratus* does not have RNA-seq data. (C) Relative gene expression in the three species measured in the current paper by RNA-seq for *P. miniata* (orange curves), in [17] by RNA-seq for *P. lividus* (purple curves) and in [23] by nanostring for *S. purpuratus* (black curves). Error bars indicate standard deviation. To obtain relative expression levels for each species we divide the level at each time point in the maximal mRNA level measured for this species in this time interval; so 1 is the maximal expression in this time interval.

The models of the gene regulatory networks that control the development of the sea urchin and those that control cell fate specification in the sea star are the state of the art in the field [3–11]. The endodermal and ectodermal gene regulatory networks show high levels of conservation between the sea urchin and the sea star in agreement with the overall conserved morphology of these two germ layers [6, 9, 11]. Surprisingly, most of the transcription factors active in the skeletogenic and the pigment mesodermal lineages are also expressed in the sea star mesoderm [7, 8]. Furthermore, the expression dynamics of key endodermal, mesodermal and ectodermal regulatory genes were compared between the Mediterranean sea urchin, *Paracentrotus lividus* (*P. lividus*) and the sea star, *Patiria miniata* (*P. miniata*) [13]. Despite the evolutionary divergence of these two species, an impressive level of conservation of regulatory gene expression in all the embryonic territories was observed. This could suggest that novel mesodermal lineages diverged from an ancestral mesoderm through only a few regulatory changes that drove major changes in downstream gene expression and embryonic morphology [7, 8].

Insight on the genome-wide changes in developmental gene expression can be gained from comparative transcriptome studies of related species at equivalent developmental stages [14–17]. We previously investigated different aspects of interspecies conservation of gene expression between *P. lividus* and the pacific sea urchin, *Strongylocentrotus purpuratus* (*S. purpuratus*) [16, 17]. These species diverged from their common ancestor about 40 million years ago and have a highly similar embryonic morphology (Fig. 1A). We observed high conservation of gene expression dynamics of both developmental genes (expressed differentially and regulate specific embryonic lineages) and housekeeping genes (expressed in all the cells throughout developmental time) [17]. Yet, the interspecies correlations of the expression levels of these two sets of genes show distinct behaviors.

The interspecies correlations of the expression levels of developmental genes are high between morphologically similar stages in the two sea urchins, and decrease sharply between diverse developmental stages, resulting with highly diagonal correlation matrices [17]. The correlations peak at mid-development, at the onset of gastrulation, in agreement with the hourglass model of developmental conservation [17–19]. Conversely, the interspecies correlations of housekeeping gene expression increase with developmental time and are high between all post-hatching time points, resulting with distinct off-diagonal elements of the correlation matrix [17]. This indicates that the ratio between the expression levels of sets of housekeeping genes is conserved and maintained throughout embryogenesis, irrespective of specific developmental stages. Interestingly, another situation where the interspecies correlation matrix is not diagonal was observed for highly diverged species that belong to different phyla [20]. In these large evolutionary distances, the interspecies correlation has an opposite hourglass behavior with high correlations between all early time points and between all late time points [20]. This suggests that the off-diagonal elements of the interspecies correlation matrix might contain information about the relationship between the conservation of gene expression and the conservation of morphology.

Apparent differences in the conservation patterns between developmental and housekeeping genes were observed in other studies of closely related species [18] and were a reason to exclude housekeeping genes from comparative studies of developing embryos [19]. Yet, embryogenesis progression depends on the dynamic expression of housekeeping genes that is highly conserved between closely related species [17]. Here we aim to decipher how the patterns of gene-expression conservation change between closely to distantly related species using the sea star and the sea urchin as our model systems. To this end, we generated and analyzed *de-novo* developmental quantitative transcriptomes of the sea star, *P. miniata*, and compared them with the published developmental transcriptomes of *P. lividus* [17] and *S. purpuratus* [21] at equivalent developmental stages (Fig. 1B). Our studies illuminate how the interspecies conservation of gene expression levels for different functional classes and cell lineages change with evolutionary distance.

## Results

### Developmental transcriptomes of the sea star, *P. miniata*

To study the transcriptional profiles of the sea star species, *P. miniata*, and compare them to those of the sea urchin species, *S. purpuratus* and *P. lividus* we collected embryos at eight developmental stages matching to those studied the sea urchin species [17, 21, 22], from fertilized egg to late gastrula stage (Fig. 1B). Details on reference transcriptome assembly, quantification and annotations are provided in the Materials and Methods section. Quantification and annotations of all identified *P. miniata* transcripts are provided in Table S2. The expression profiles of selected regulatory genes show mild heterochronies and overall similarity between *P. miniata* (RNA-seq, current study) and the sea urchin species, *S. purpuratus* (nanostring, [23]) and *P. lividus* (RNA-seq, [17]), in agreement with our previous study (Fig. 1C, [12]).

### Identification of 1:1:1 orthologues gene set and quantitative data access

To compare developmental gene expression between the sea star, *P. miniata*, and the two sea urchins, *P. lividus* and *S. purpuratus* we identified 8735 1:1:1 putative homologous genes, as described in the Materials and Methods section. Quantification and annotations of all these homologous genes in *P. miniata*, *P. lividus* and in *S. purpuratus* are provided in Table S3 (based on [17] for *P. lividus* and on [21, 22] *for S. purpuratus*). Within this set, 6593 genes were expressed in the three species, 1093 genes are expressed only in the two sea urchin species, 187 genes were expressed only in the sea star, *P. miniata*, and the sea urchin, *S. purpuratus*, and 430 genes are expressed only in *P. miniata* and *P. lividus* (Fig. 2A). We looked for enrichment of gene ontology (GO) terms within these different gene sets but did not identify enrichment of specific developmental processes (GOseq [24] with *S. purpuratus* annotations, Fig. S1 and Table S4).

**Figure 2.**
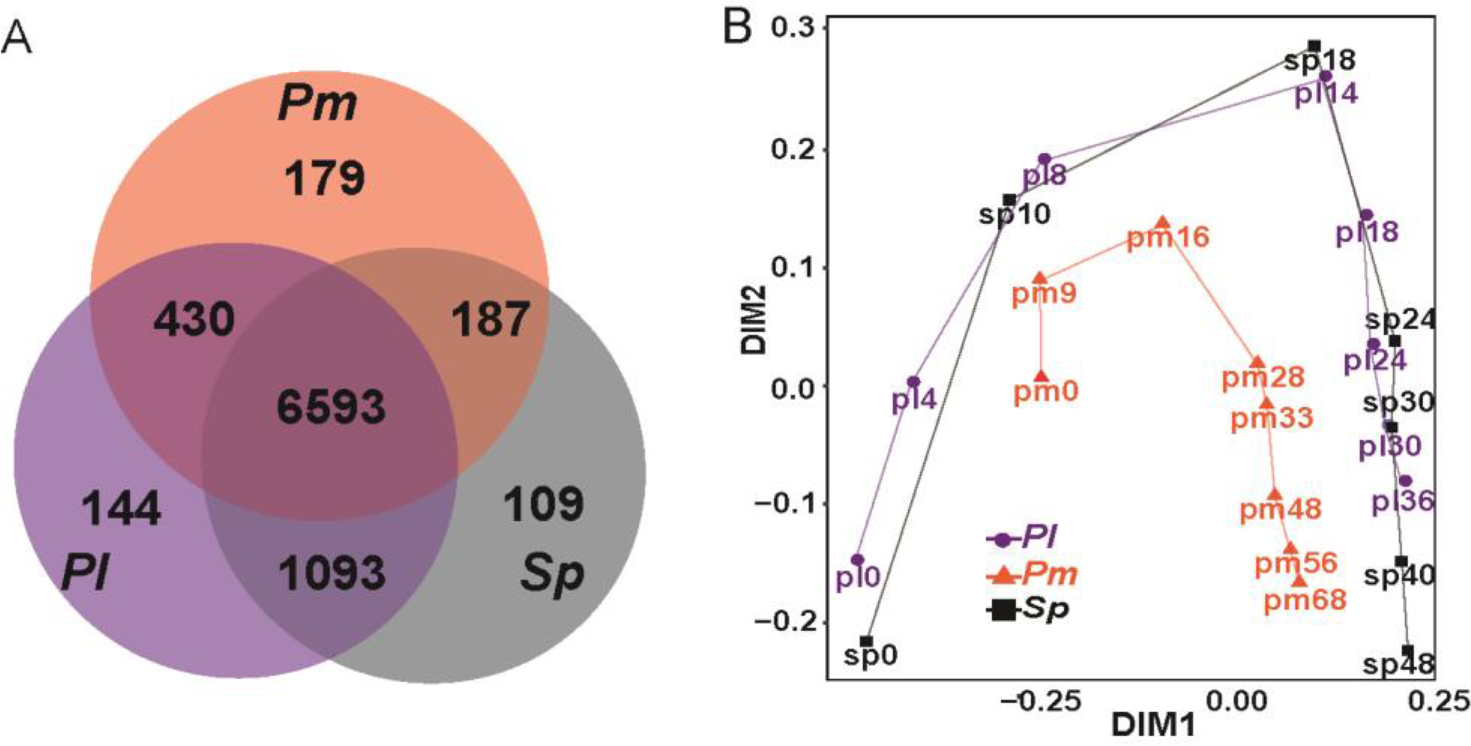
Venn diagram and NMDS analysis of 1:1:1 orthologues genes. (A) Venn diagram showing the number of 1:1:1 orthologues genes expressed in all three species, in two of the species or only in one species. (B) First two principal components of expression variation (NMDS) between different developmental time points in *P. miniata* (orange), *P. lividus* (purple) and *S. purpuratus* (black).

We uploaded the data of the 1:1:1 homologous genes expressed in the three species to Echinobase where they are available through gene search at www.echinobase.org/shiny/quantdevPm [25]. In this Shiny web application [26], genes can be searched either by their name or by their *P. miniata*, *S. purpuratus* or *P. lividus* transcript identification number. The application returns our quantitative measurements of gene expression for each stage of *P. miniata*, plots of transcript expression level, mRNA sequences, a link to the corresponding records in Echinobase [27] and links to the loci in the *S. purpuratus* and *P. miniata* genome browsers. This will hopefully be a useful resource for the community. Further analyses of the 6593 1:1:1 homologous genes expressed in all three species are described below.

### Similarity in gene expression profiles depends on developmental stage and evolutionary distance

To identify the most similar stages in terms of gene expression within and between species, we performed a Non-metric multidimensional scaling analysis (NMDS, Fig. 2B). *S. purpuratus* RNA-seq data does not include the early development time point equivalent to the *P. miniata* 9hpf and *P. lividus* 4hpf [21]. The NMDS maps the equivalent developmental time points of the two sea urchin species closely to each other (purple and black tracks), while the sea star samples are relatively distinct (orange track). This is in agreement with the relative evolutionary distances between the three species (Fig. 1A). However, the developmental progression in all three species is along a similar trajectory in the NMDS two dimensional space, which probably reflects the resemblance in the overall morphology between these three echinoderm species. Overall, the NMDS indicates that major sources of variation in these data sets are developmental progression and evolutionary distance.

### Interspecies correlations decrease and the pattern becomes less diagonal with evolutionary distance

We wanted to study how the pattern of the interspecies correlations of gene expression changes with evolutionary distance for different classes of genes. To be able to compare the interspecies correlations between the three species we included only time points that had data for all species. Explicitly, we excluded *P. lividus* 4hpf and *P. miniata* 9hpf that do not have an equivalent time point in the *S. purpuratus* data. We calculated the Pearson correlations of gene expression levels between *P. lividus* and *S. purpuratus* and between *P. lividus* and *P. miniata* for subsets of homologous genes with specific developmental, housekeeping or metabolic function, according to their GO terms (Fig. 3). As expected, the interspecies correlation patterns differ in strength and diagonality between different functional classes for both *Pl-Sp* and *Pl-Pm* comparison.

**Figure 3.**
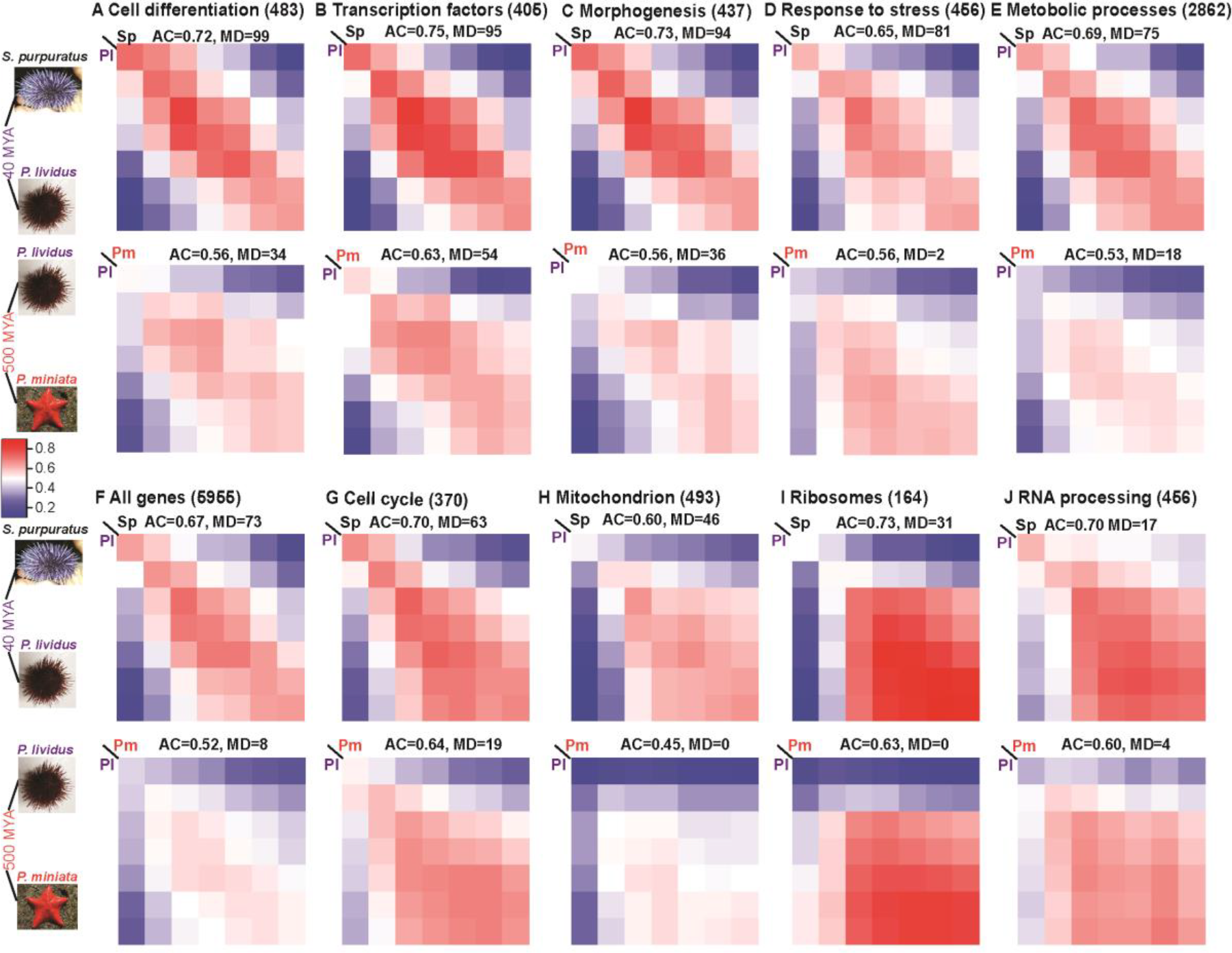
Interspecies Pearson correlations for different GO terms, ordered by the level of matrix diagonality (MD) of *Pl-Sp* matrices. In each panel, from A to J, we present the Pearson correlation of the expression levels of genes with specific GO term between different developmental stages in the three species. Upper matrix in each panel shows the Pearson correlation between the two sea urchins (*P. lividus* and *S. purpuratus*) and the bottom matrix is the Pearson correlation between the sea star, *P. miniata* and the sea urchin *P. lividus*. These matrices include the seven developmental points that have RNA-seq data in all species (Fig. 1B, excluding 9hpf in *P. miniata* and 4hpf in *P. lividus*). In each panel we indicate the GO term tested, the number of genes in each set, the average correlation strength in the diagonal (AC) and the matrix diagonality (MD), see text for explanation. Color scale is similar for all graphs and given at the middle of the figure. F, shows the Pearson interspecies correlation for all 1:1:1 genes.

For a better assessment of the correlation pattern we sought to quantify these two distinct properties: the correlation strength between equivalent developmental stages and the diagonality of the matrix. As a measure of the correlation strength between morphologically similar time points, we defined the parameter **AC = Average Correlation, which is the average of the diagonal elements of the correlation matrix**. That is, AC=1 corresponds to perfect correlation throughout all equivalent developmental stages and AC=0 corresponds to no correlation. Evidently, AC is not a measure of the diagonality of the matrix as it doesn’t consider the off-diagonal terms. To quantify the diagonality of the correlation matrices we used a statistical test we developed before to assess whether the interspecies correlation pattern is significantly close to a diagonal matrix [17]. Briefly, the parameter, **matrix diagonality (MD), indicates how many times within 100 subsamples of the tested set of genes, the interspecies correlation matrix was significantly close to a diagonal matrix compared to a random matrix**. Hence, MD=100 is the highest diagonality score and 0 is the lowest (see materials and methods and [17] for explanation, and Table S5 for results. MD is equivalent to ‘count significant’, or CS, in [17]). Importantly, AC will be high for any matrix that has high diagonal elements, but MD will be high only when the off-diagonal matrix elements are much lower than the diagonal, and low otherwise. We ordered the correlation matrices according to their MD value in *Sp-Pl*, from the most diagonal (Cell differentiation, MD=99) to the least (RNA processing, MD=17). This ordering clearly demonstrates the high diagonality of the developmental genes (Fig. 3A-C) vs. the block patterns of the housekeeping genes (Fig. 3G-J) and how the diagonality of the correlation pattern of all genes combined is in between (Fig. 3F). Both the average correlation and the matrix diagonality are lower between the sea urchin and the sea star than between the two sea urchins, that is, both parameters decrease with evolutionary distance (compare bottom to upper panels in Fig. 3A-I).

Low correlation matrix diagonality indicates that the off-diagonal elements have a similar strength to the diagonal elements. This means that the relationship between the expression levels of a set of genes is maintained not only between morphologically similar developmental stages, but throughout development. In the case of housekeeping genes that form complex structures, like ribosomal proteins, this could be explained by stoichiometric ratio requirements between different proteins that form the structure. With increasing evolutionary distance and morphological divergence there is a reduction in matrix diagonality for all sets of genes (Fig. 3). Possibly, the dominant constraints in large evolutionary distances are the cellular requirement for different levels of transcripts that have different functions. These constraints are cellular and not related to specific morphogenetic processes, which might explain the even correlation strength between different developmental stages and hence, the lower diagonality at large evolutionary distances.

### The interspecies correlation vary between different cell lineages

Our analysis is based on RNA-seq on whole embryos, yet we wanted to see if gene sets enriched in a specific cell lineage show a difference in their interspecies correlation pattern. In a previous work conducted in the sea urchin *S. purpuratus* embryos, the cells of six distinct embryonic lineages were isolated based on cell-specific GFP reporter expression. Gene expression levels in the isolated cells were studied and compared to gene expression levels in the rest of the embryo by RNA-seq [28]. This analysis identified genes that their expression is enriched in the specific lineages at the time when the isolation was done [28]. We calculated the Pearson interspecies correlations for the subsets of genes enriched in each of the six sea urchin lineages, between the two sea urchins and between the Mediterranean sea urchin and the sea star (Fig. 4). The sea urchin cell lineages at the time of cell isolation are illustrated above the correlation patterns and the developmental stage is indicated as a black square within the *Pl-Sp* correlation matrices.

**Figure 4.**
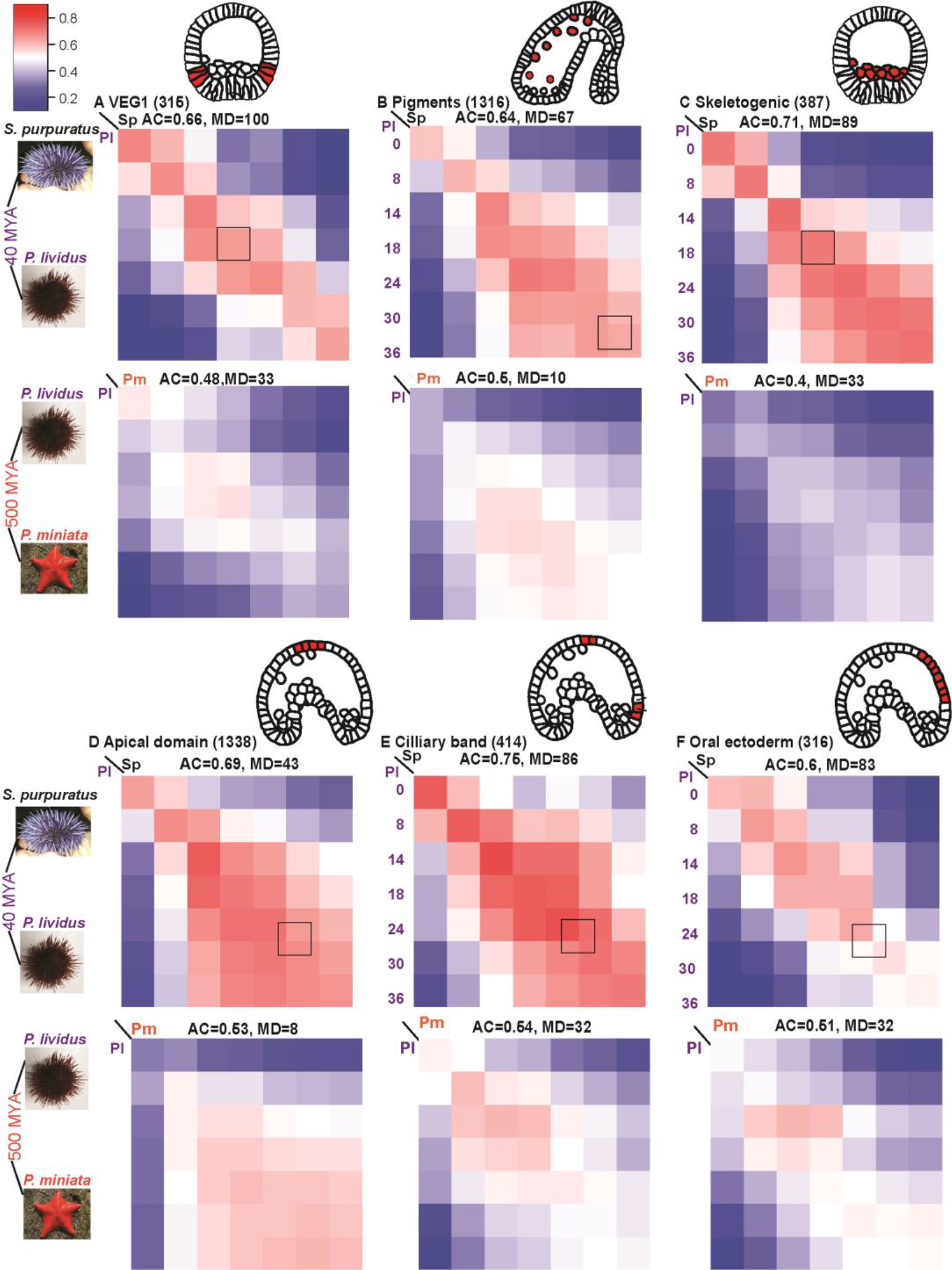
Interspecies Pearson correlations in gene expression for genes enriched in specific sea urchin lineages. A-F, each panel shows the interspecies Pearson correlation between the developmental stages in the three species for genes that their expression enriched at specific time point in a particular cell lineage in the sea urchin*, S. purpuratus* according to [28]. The cell lineages at the time where the enrichment was observed are illustrated by the embryo diagrams above the relevant correlation pattern[28], the time point is also marked in a black square in each *Pl-Sp* panel. In each panel we indicate lineage were these genes are enriched, the number of genes in each set, the average correlation strength in the diagonal (AC) and the matrix diagonality (MD), see text for explanation. Color scale is similar for all graphs and given at the top.

Both the matrix diagonality and average correlation strength vary between the different cell types and decrease with evolutionary distance. Interestingly, the genes enriched in sea urchin pigment cells, a lineage that is lacking in the sea star, show similar *Pl-Pm* average correlation strength compared to the matrices of the conserved lineages between the sea urchin and the sea star (*Pl-Sp* AC=0.64 and *Pl-Pm* AC=0.5, Fig. 4B). As mentioned above, the regulatory state in the mesoderm of the sea star and the sea urchin are quite similar [7]. Possibly, there is also similarity in the downstream genes active in sea star blastocoelar cells and sea urchin pigment cells, as both lineages have hematopoietic function [29]. On the other hand, the sharpest decrease in *Pl-Pm* average correlation strength compared to *Pl-Sp* is for the genes enriched in the sea urchin skeletogenic cells, another lineage that is absent in the sea star (*Pl-Sp* AC=0.71 while *Pl-Pm* AC=0.4, Fig. 4C). Interestingly, the *Pl-Pm* matrix diagonality of these genes is comparable to the *Pl-Pm* matrix diagonality of the conserved lineages (*Pl-Pm* MD=0.33, Fig. 4C). It is important to note that key sea urchin skeletogenic matrix proteins were not found in the sea star skeleton [30] and the genes encoding them were not found in the sea star genome. Therefore these key skeletogenic genes are missing from our 1:1:1 homologous genes that include only genes that are common to the three species. Thus, we are probably underestimating the differences in skeletogenic and mesodermal gene expression between the sea urchin and the sea star, which might explain the relatively high diagonality for the skeletogenic gene correlation matrix. Overall, genes enriched in different cells lineages differ in their correlation pattern even between the closely related sea urchins and show distinct differences in the correlation pattern with evolutionary distance.

### Matrix diagonality and correlation strength describe different properties of gene expression conservation

Both the average correlation strength and the matrix diagonality depend on gene function, cell lineage and decrease with evolutionary distance (Figs. 3, 4). When we plot, separately, the average correlation and the matrix diagonality for different GO terms and cell lineages, in decreasing strength in *Pl-Pm* we see two distinct orders (Fig. 5A and B). While the matrix diagonality separates clearly the developmental genes with high diagonality from the housekeeping genes with low diagonality, the average correlation does not show this distinction (Fig. 5A, B). Moreover, the matrix diagonality and the average correlation strength change independently of each other in both *Pl-Sp* and *Pl-Pm* correlation matrices (Fig. 5C). On the other hand, the average correlations in *Pl-Sp* matrices seem to correspond to the average correlation in *Pl-Pm* (Fig. 5D), and the same is true for the matrix diagonality (Fig. 5E).

**Figure 5.**
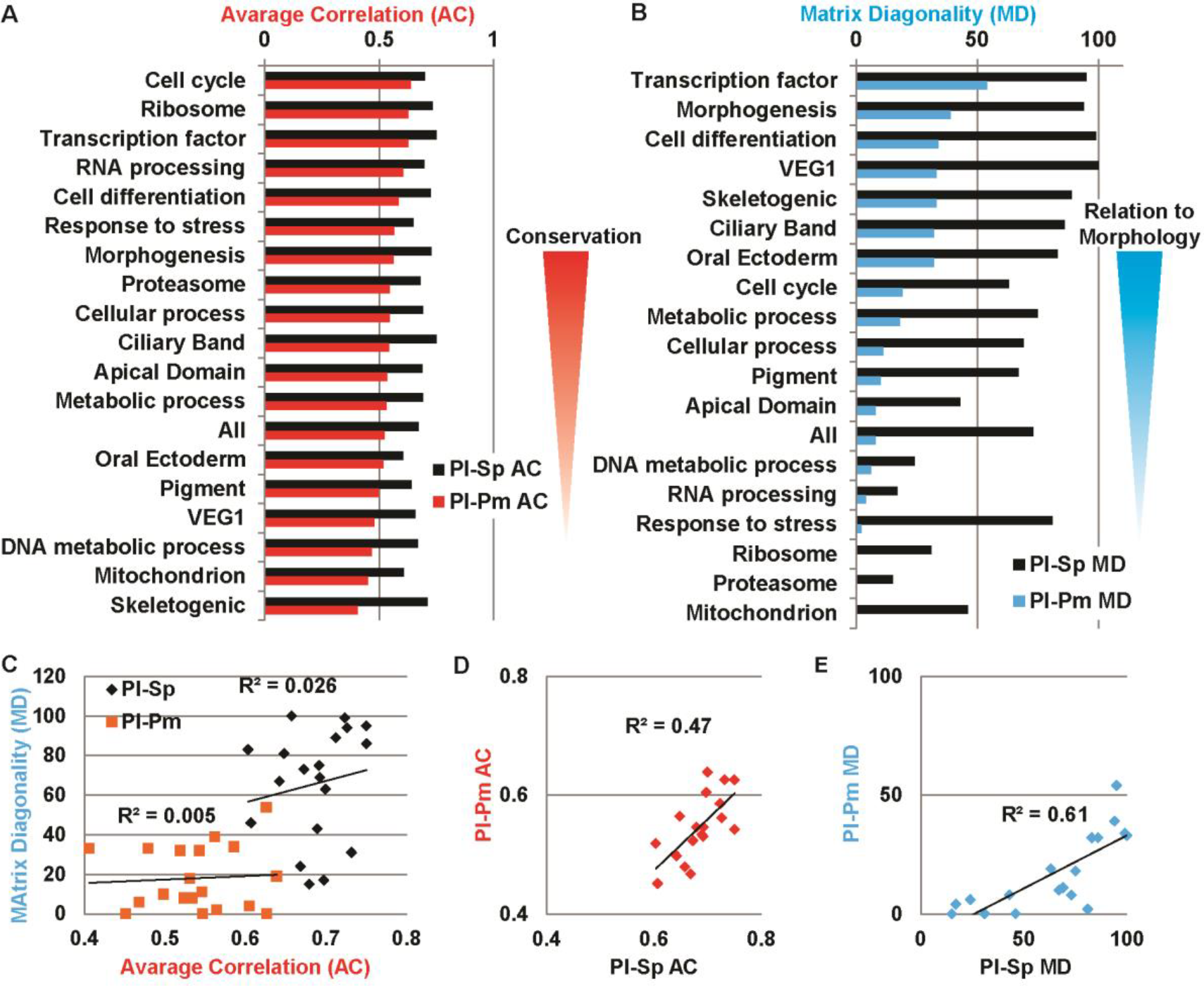
The average correlation strength and the matrix diagonality are independent parameters that reflect different properties of expression conservation. A, the average correlation strength in the diagonal (AC) between *Pl-Sp* (black bars) and *Pl-Pm* (red bars) in receding order of *Pl-Pm* correlation strength. B, Matrix diagonality (MD) of the interspecies correlations between *Pl-Sp* (black bars) and *Pl-Pm* (cyan bars) in receding order of *Pl-Pm* matrix diagonality. C, the matrix diagonality changes independently of the average correlation for both *Pl-Sp* (black dots) and *Pl-Pm* (orange dots). D, the interspecies average correlation between *S. purpuratus* and *P. lividus* corresponds to the interspecies average correlation between *P. lividus* and *P. miniata* (excluding skeletogenic genes, R pearson = 0.68). E, the matrix diagonality of the interspecies correlations between *S. purpuratus* and *P. lividus* relates to the matrix diagonality between *P. lividus* and *P. miniata* (excluding skeletogenic genes, R Pearson = 0.78).

These analyses imply that these two parameters describe different properties of the conservation in gene expression, as we illustrate in Fig. 6. Apparently, the average correlation strength is a measure of the conservation in gene expression levels: the higher it is, the more the gene-set is constrained against expression change. The matrix diagonality on the other hand, seems to reflect the link between the expression of a gene-set and morphological conservation: the more diagonal is the correlation pattern of a gene set, the more related is the gene expression conservation within the set to morphological similarity (Fig. 6). For example, the correlation matrices of transcription factors and cell differentiation genes show strong correlations and high diagonality even between the sea urchin and the sea star (Relatively high AC and MD, Fig. 3A,B, 6). Conversely, ribosomal gene expression is highly conserved (high AC) but this conservation is not related to morphological conservation (Low MD, Fig. 3I, 5A,B, 6). Overall, the average correlation strength and matrix diagonality seem to carry complementary information about the relationship between the conservation of gene expression and morphological conservation at varying evolutionary distances (Fig. 6).

**Figure 6.**
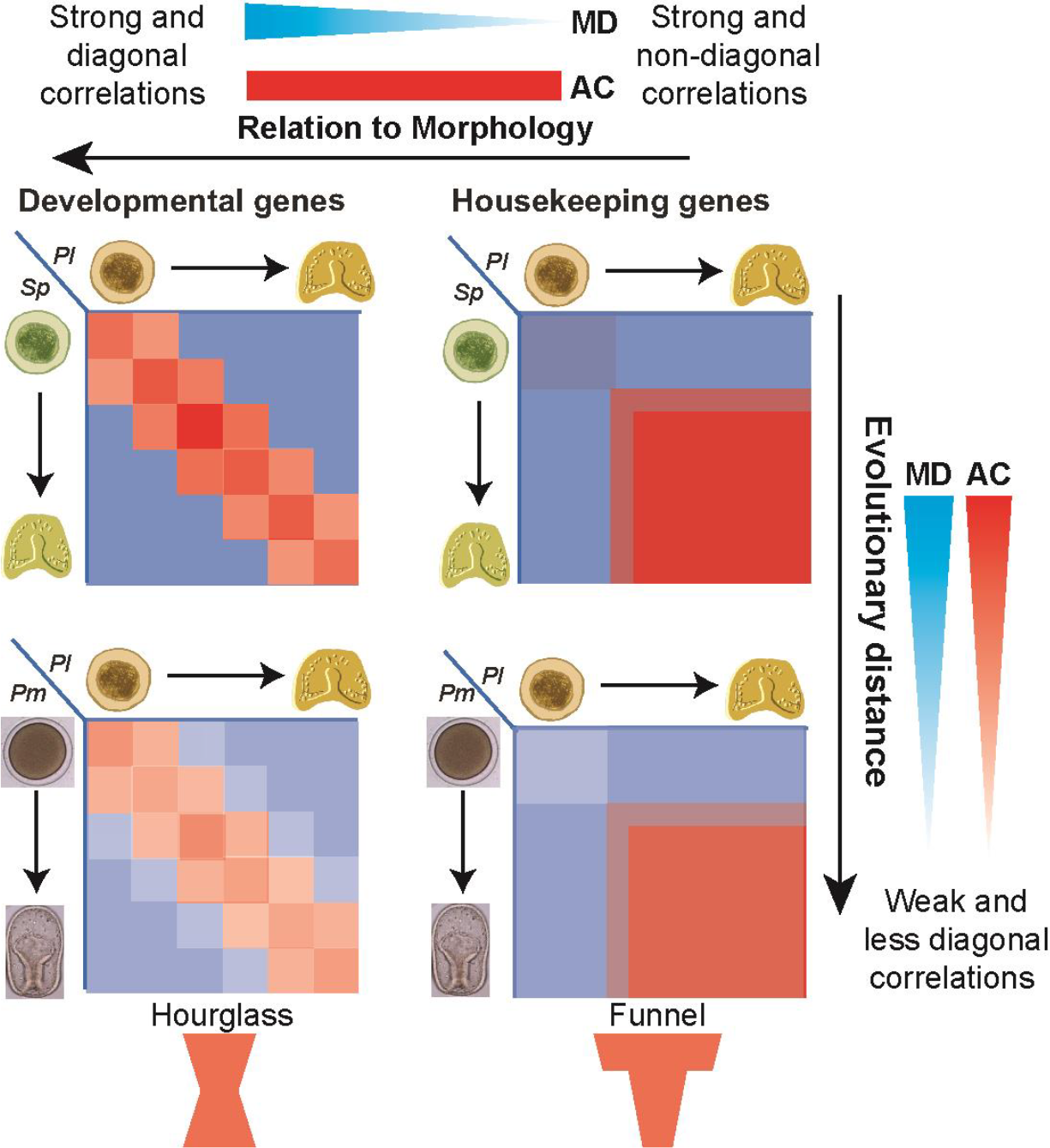
The correlation matrix diagonality, MD, reflects how the dominance between cellular and developmental constraints changes with evolutionary distance for different functional classes. Illustration of typical interspecies correlation matrices of developmental control genes and housekeeping genes between closely related and further diverged (lower species (upper and lower panels, respectively). With increasing evolutionary distance, that is, between the sea urchin and the sea star, the average correlation and the diagonality decrease for all gene sets but the diagonality of developmental control genes is least affected and they that still maintain the hourglass pattern (lower left panel). On the other hand, the interspecies correlation matrices of housekeeping genes are strong and non-diagonal even between the two sea urchins and remain non-diagonal between the sea urchin and the sea star (right panels).

## Discussion

In this paper we generated the developmental transcriptomes of the pacific sea star, *P. miniata*, and studied them in comparison with the published developmental transcriptomes of two sea urchin species, *P. lividus* [17] and *S. purpuratus* [21]. We generated a web application where the *P. miniata* time courses and sequences can be publically viewed to facilitate the common use of this data [25]. We studied the interspecies correlation patterns of different gene sets including, housekeeping, developmental and metabolic genes (Fig. 3), as well as genes that are enriched in a specific cell lineage in the sea urchin embryo (Fig. 4). We defined two parameters that describe different properties of the conservation strength: the average correlation strength in the diagonal, AC, and the matrix diagonality, MD. We noticed that these parameters vary independently between different functional groups and cell types and decrease with evolutionary distance, possibly reflecting different constraints on gene expression (Fig. 5). The correlation strength is an indication of the evolutionary constraint on gene expression level while the matrix diagonality seems to reflect the relationship between the gene set and morphological similarity. As we suggested previously, parallel embryonic transcriptional programs might be responsible for different aspects of embryo development and evolve under distinct constraint [17], as can be inferred from analyzing these two parameters.

Previous studies have shown that housekeeping genes and tissue specific genes have different chromatin structures [31] and distinct core promoters [32]. Furthermore, Enhancers of developmental genes were shown to be de-methylated during the vertebrates’ phylotypic period, suggesting another unique epigenetic regulation of this set of genes [33]. These epigenetic and *cis*-regulatory differences could underlie the separation of the regulation of developmental and housekeeping gene expression, leading to dissimilar evolutionary conservation patterns of these two functionally distinct gene sets.

We observed a clear reduction of the matrix diagonality with increasing evolutionary distance for all sets of genes, including developmental genes and transcription factors (Fig. 3–6). This is in agreement with the morphological divergence between the sea urchin and the sea star. As mentioned above, a recent study had shown that the correlation matrices for all homologous genes between 10 different phyla are strictly not-diagonal [20]. At this large evolutionary distance and extreme morphological divergence the only constraint that remains seem to be the cellular requirements for differential transcript abundance, which are unrelated to morphological similarity. Therefore the opposite hourglass pattern found for the diagonal elements of the interspecies correlation matrices might not be indicative for morphological divergence at the phylotypic stage. Overall, to better estimate the relationship between the conservation of gene expression levels to morphological conservation, both correlation strength and correlation matrix diagonality should be assessed and the focus should be on developmental control genes.

## Materials and Methods

### Sea star embryo cultures and RNA extraction

Adult *P. miniata* sea stars were obtained in Long Beach, California, from Peter Halmay. Embryos were cultured at 15°C in artificial sea water. Total RNA was extracted using Qiagen mini RNeasy kit from thousands of embryos. RNA samples were collected from eight embryonic stages, from fertilized egg to late gastrula stage (Fig. 1B). For each embryonic stage, three biological replicates from three different sets of parents were processed, except for the last time point for which only two biological replicates were samples (23 samples in all). To match *P. miniata* time points to those of the published *S. purpuratus* and *P. lividus* transcriptomes [17, 22] we used the linear ratio between the developmental rates of these species found in [12, 34]. The corresponding time points in both species are presented in Fig. 1B. The time points, 9 hour post fertilization (hpf) in *P. miniata* and 4hpf in *P. lividus* do not have a comparable time point in *S. purpuratus* data. RNA quantity was measured by nanodrop and quality was verified using bioanalyzer.

### Transcriptome assembly, annotations and quantification

#### Transcriptome assembly, annotations and quantification

##### RNA-Seq preparation

Library preparation was done at the Israel National Center for Personalized Medicine (INCPM) using their RNA-seq protocol. Briefly, polyA fraction (mRNA) was purified from 500 ng of total RNA per developmental time point following by fragmentation and generation of double stranded cDNA. Then, end repair, A base addition, adapter ligation and PCR amplification steps were performed. Libraries were evaluated by Qubit and TapeStation. Sequencing libraries were constructed with barcodes to allow multiplexing of 47 samples in one lane. On average, 20 million single-end 60-bp reads were sequenced per sample on Illumina HiSeq 2500 V4 using four lanes.

##### RNA-Seq datasets used

Four datasets were used: (1) newly sequenced *P. miniata* Single End (SE) reads of 23 transcriptome samples generated as explained above; (2) publically available Paired End (PE) reads of *P. miniata* from different developmental stages, testis and ovaries, accessions: SRR6054712, SRR5986254, SRR2454338, SRR1138705, SRR573705-SRR573710 and SRR573675; (3) publically available SE reads of *P. lividus* transcriptome samples from eight developmental stages (NCBI project PRJNA375820, [17]); (4) publically available PE reads of seven *S. purpuratus* transcriptome samples (NCBI project PRJNA81157, [22]).

##### RNA-seq quality filtering

RNA-Seq reads from the above datasets were adapter-trimmed using cutadapt 1.15 (https://cutadapt.readthedocs.io/), then low-quality regions were removed with Trimmomatic 0.3 [35], and further visually inspected in fastq-screen (www.bioinformatics.babraham.ac.uk).

##### *P. miniata* transcriptome assembly

While *P. miniata* genome based gene predictions are available for genome assembly v.2 (echinobase.org/Echinobase), many gene sequences are fragmented or duplicated within the genome, and therefore we decided to generate a reference transcriptome *de-novo*. Accordingly, *P. miniata* RNA-Seq reads, based on the available PE and SE data, were assembled using Trinity 2.4 [36, 37] with the same Trinity parameters as before we used for *P. lividus* in [17]. Trinity produced 1,610,829 contigs (the Trinity equivalent of transcripts), within 679,326 Trinity gene groups. Transciptome completeness was tested using BUSCO [38], and by comparing the contigs to genome-based protein annotations *P. miniata* v.2.0 and *S. purpuratus* v.3.1 (WHL22), (echinobase.org/Echinobase).

##### *P. miniata* data availability

Illumina short read sequences generated in this study was submitted to the NCBI Sequence Read Archive (SRA) (http://www.ncbi.nlm.nih.gov/sra), under bio-project PRJNA522463 (http://www.ncbi.nlm.nih.gov/bioproject/522463). Fastq read accessions: SRR8580044 - SRR8580066, assembled *P. miniara* transcriptome accession: SAMN10967027.

##### Transcriptome homology

We searched for homologous genes in the *P. miniata* transcriptome, *P. lividus* transcriptome*, S. purpuratus* genome. Since the *S. purpuratus* genome-based gene predictions dataset currently includes the most non-redundant and complete data among the three datasets, it was used as a reference dataset. Accordingly, the largest isoforms of new *P. miniata* transcriptome (see below), and the publicly available *P. lividus* transcriptome [17], were both compare to the *S.purpuratus* genome-based predicted protein annotations (Echinobase v.3.1), using CRB-BLAST [39]. CRB-BLAST reports relationships of 1:1 (Reciprocal hits), and 2:1 matches (when Blastx and tBlastn results are not reciprocal), where only matches with e-values below a conditional threshold are reported. From the 2:1 cases, we selected the query-target pair with the lowest e-value. Using CRB-BLAST, 11,291 and 12,720 *P. miniata* and *P. lividus* Trinity genes, were identified as homologous to *S. purpuratus* proteins, respectively. We considered *P. miniata* and *P. lividus* query genes that share the same *S. purpuratus* target gene, as homologous. As the gene expression analysis shows (see next sections), 8,735 homologous genes are expressed in at least one of the three tested species during development, and 6,593 are expressed in all the three.

##### Gene-level transcripts abundance

For *P. miniata* and *P. lividus* transcriptomes, transcripts abundance was estimated using kallisto-0.44.0 [40], and a further quantification at the gene-level, and read-count level, was done using tximport [41] on R3.4.2. Expression analysis at read-count level was conducted at gene-level in Deseq2 [42]. *S. purpuratus* PE reads, of 7 developmental samples were mapped to the *S. Purpuratus 3.1* genome assembly, using STAR v2.4.2a [43], quantitated in Htseq-count v2.7 [44], and analyzed in Deseq2 at read-count level. For *P. miniata* data, only contigs mapped to the Echinobase v.2 *P. miniata* proteins were considered. Prior to DEseq2 read count standard normalization and expression analysis, genes with <1 CPM (Count Per Million) were removed. Overall, most input reads were mapped and quantified, as further detailed in Table S1. Samples were clustered using Non-metric multi-dimentional scaling (NMDS) ordination in Vegan (https://cran.r-project.org/web/packages/vegan/index.html), based on log10 transformed FPKM values, and Bray-Curtis distances between samples. Since the NMDS results indicate that all *P. miniata* and *P. lividus* samples are affected by the ‘batch’ factor (see Table S1), we removed the estimated effect of this factor on FPKMs (Fragment per Kilobase Million) values using “removeBatchEffect” function in EdgeR [45], in order to obtain corrected FPKM values. Quantification and annotations of 34,307 identified *P. miniata* transcripts with FPKM>3 in at least one time point, are provided in Table S2. NMDS analysis of the biological replicates of all time points in *P. miniata* show that similar time points at different biological replicate map together indicating high reproducibility of our gene expression analysis (Fig. S2). Comparison between our RNA-seq quantification of gene expression and previous QPCR quantification [12] for a sub-set of genes show high agreement between the two measurements (Fig. S3). Quantification and annotations of the 8,735 identified *P. miniata, P. lividus and S. purpuratus* 1:1:1 homologous genes are provided in Table S3 and are publicly available through gene search in Ehinobase at www.echinobase.org/shiny/quantdevPm.

##### Gene Ontology functional enrichment analysis

Functional enrichment analysis was conducted using TopGo in R3.4.2 (bioconductor.org). A custom GOseq GO database was built using the publically available Blast2Go *S. purpuratus v3.1* (WHL22) version (http://www.echinobase.org/Echinobase/rnaseq/download/blast2go-whl.annot.txt.gz).

##### Pearson correlations for subsets of genes

The interspecies Pearson correlations for different sets of genes presented in Figs. 3 and 4 were calculated using R3.4.2. For the analysis in Fig. 4, we selected genes that have 1:1:1 homologs in *P. miniata* and *P. lividus* to the *S. purpuratus* genes that are enriched in a specific *S. purpuratus* cell lineage with p-value<0.05, based on [28].

##### Cross species analysis of matrix diagonality (MD)

We used a statistical test described in detail in [17]. Shorty, the main goal of this procedure is to test the probability that a set of homologous genes from two species, *S*_1_ and *S*_2_, show the most similar expression patterns in equivalent developmental times, namely: is the interspecies correlation pattern significantly close to a diagonal matrix? Here, *S*_1_ and *S*_2_ represent *P. miniata* vs. *P. lividus*, or *P. lividus* vs. *S. purpuratus*. This test was conducted using all homologous genes, and for specific subsets of genes belonging to specific GO categories as well as for genes enriched in specific cell lineage [28]. Here, only samples from the *n*=5 late embryonic stages were used, and the mean of FPKM values of all samples belonging to the same stages were taken. First, *n*_*s1*_ by *n*_*s2*_ matrix of Pearson correlations, *C*, was produced, where *n*_*s1*_ by *n*_*s2*_ are the number of time point measurements in each species (here *n*_*s1*_ =*n*_*s2*_ =5). Each of the *C* matrix positions, *C*_*ij*_, represents a Pearson’s correlation value calculated based on *n*_*g*_ FPKM values, between stage *i* in one species and stage j in the other, where *n*_*g*_ is the count of homologous gene pairs tested. Next, the *C* matrix was compared to an “ideal” time-dependent correlation matrix *I*, in which a correlation of 1 exists along the diagonal line, and 0 in other positions, to obtain a diagonality measure, *d*. Overall, *n*_*g*_ =50 genes were resampled *n*_*resamp*_ =100 times, and for each resampling-iteration, a permutation test based on value *d* was applied using *n*_*perm*_ =1000 permutations. Accordingly, each of the above *n*_*resamp*_ =100 permutation tests indicate the probability that *S*_1_ and *S*_*2*_ show non-random diagonality, on a subset of n_g_=50 genes. Then, the count of permutation tests indicating a significant similarity to the ideal diagonal matrix, out of the *n*_*resamp*_ =100 subsamples is used for estimating diagonality of the correlation matrix (matrix diagonality = MD). Our results are presented in Table S5 and within Figs. 3–5.

## Acknowledgements

We thank Assaf Malik from the bioinformatics service unit at the University of Haifa for the analyses of the transcriptome data. We thank the Genomics unit at the Grand Israel National Center for Personalized Medicine (G-INCPM) for their assistance with library preparation and sequencing. We thank Paola Oliveri for the idea to study lineage specific genes. This work was supported by the Binational Science Foundation grant number 2015031 to SBD and VFH and the Israel Science Foundation grants number 2304/15 (ISF-INCPM) to SBD, the National Science Foundation grants IOS 1557431 and MCB 1715721 to VFH and the National Institute of Health grant P41HD071837 to VFH.

## References

1. Peter IS, Davidson EH. Genomic Control Process: Development and Evolution. New York: Academic Press; 2015.

2. Raff RA, Byrne M. The active evolutionary lives of echinoderm larvae. Heredity. 2006;97(3):244–52. Epub 2006/07/20. doi: 10.1038/sj.hdy.6800866. PubMed PMID: 16850040.

3. Oliveri P, Tu Q, Davidson EH. Global regulatory logic for specification of an embryonic cell lineage. Proc Natl Acad Sci U S A. 2008;105(16):5955–62. Epub 2008/04/17. doi: 10.1073/pnas.0711220105. PubMed PMID: 18413610; PubMed Central PMCID: PMC2329687.

4. Ben-Tabou de-Leon S, Su YH, Lin KT, Li E, Davidson EH. Gene regulatory control in the sea urchin aboral ectoderm: Spatial initiation, signaling inputs, and cell fate lockdown. Dev Biol. 2012. Epub 2012/12/06. doi: 10.1016/j.ydbio.2012.11.013. PubMed PMID: 23211652.

5. Li E, Materna SC, Davidson EH. Direct and indirect control of oral ectoderm regulatory gene expression by Nodal signaling in the sea urchin embryo. Dev Biol. 2012;369(2):377–85. Epub 2012/07/10. doi: 10.1016/j.ydbio.2012.06.022. PubMed PMID: 22771578; PubMed Central PMCID: PMC3423475.

6. Hinman VF, Nguyen AT, Cameron RA, Davidson EH. Developmental gene regulatory network architecture across 500 million years of echinoderm evolution. Proc Natl Acad Sci U S A. 2003;100(23):13356–61. Epub 2003/11/05. doi: 10.1073/pnas.2235868100 2235868100 [pii]. PubMed PMID: 14595011; PubMed Central PMCID: PMC263818.

7. McCauley BS, Weideman EP, Hinman VF. A conserved gene regulatory network subcircuit drives different developmental fates in the vegetal pole of highly divergent echinoderm embryos. Dev Biol. 2010;340(2):200–8. Epub 2009/11/28. doi: S0012-1606(09)01375-X [pii] 10.1016/j.ydbio.2009.11.020. PubMed PMID: 19941847.

8. McCauley BS, Wright EP, Exner C, Kitazawa C, Hinman VF. Development of an embryonic skeletogenic mesenchyme lineage in a sea cucumber reveals the trajectory of change for the evolution of novel structures in echinoderms. Evodevo. 2012;3(1):17. Epub 2012/08/11. doi: 10.1186/2041-9139-3-17. PubMed PMID: 22877149.

9. Yankura KA, Martik ML, Jennings CK, Hinman VF. Uncoupling of complex regulatory patterning during evolution of larval development in echinoderms. BMC Biol. 2010;8:143. Epub 2010/12/02. doi: 1741-7007-8-143 [pii] 10.1186/1741-7007-8-143. PubMed PMID: 21118544; PubMed Central PMCID: PMC3002323.

10. McCauley BS, Akyar E, Filliger L, Hinman VF. Expression of wnt and frizzled genes during early sea star development. Gene Expr Patterns. 2013;13(8):437–44. Epub 2013/08/01. doi: 10.1016/j.gep.2013.07.007. PubMed PMID: 23899422.

11. Yankura KA, Koechlein CS, Cryan AF, Cheatle A, Hinman VF. Gene regulatory network for neurogenesis in a sea star embryo connects broad neural specification and localized patterning. Proc Natl Acad Sci U S A. 2013;110(21):8591–6. Epub 2013/05/08. doi: 10.1073/pnas.1220903110. PubMed PMID: 23650356; PubMed Central PMCID: PMC3666678.

12. Gildor T, Hinman V, Ben-Tabou-De-Leon S. Regulatory heterochronies and loose temporal scaling between sea star and sea urchin regulatory circuits. Int J Dev Biol. 2017;61(3-4-5):347–56. doi: 10.1387/ijdb.160331sb. PubMed PMID: 28621432.

13. Cary GA, Hinman VF. Echinoderm development and evolution in the post-genomic era. Dev Biol. 2017;427(2):203–11. doi: 10.1016/j.ydbio.2017.02.003. PubMed PMID: 28185788.

14. Dylus DV, Czarkwiani A, Blowes LM, Elphick MR, Oliveri P. Developmental transcriptomics of the brittle star Amphiura filiformis reveals gene regulatory network rewiring in echinoderm larval skeleton evolution. Genome Biol. 2018;19(1):26. doi: 10.1186/s13059-018-1402-8. PubMed PMID: 29490679; PubMed Central PMCID: PMCPMC5831733.

15. Israel JW, Martik ML, Byrne M, Raff EC, Raff RA, McClay DR, et al. Comparative Developmental Transcriptomics Reveals Rewiring of a Highly Conserved Gene Regulatory Network during a Major Life History Switch in the Sea Urchin Genus Heliocidaris. PLoS Biol. 2016;14(3):e1002391. doi: 10.1371/journal.pbio.1002391. PubMed PMID: 26943850; PubMed Central PMCID: PMCPMC4778923.

16. Gildor T, Smadar BD. Comparative Studies of Gene Expression Kinetics: Methodologies and Insights on Development and Evolution. Frontiers in genetics. 2018;9:339. doi: 10.3389/fgene.2018.00339. PubMed PMID: 30186312; PubMed Central PMCID: PMCPMC6113378.

17. Malik A, Gildor T, Sher N, Layous M, Ben-Tabou de-Leon S. Parallel embryonic transcriptional programs evolve under distinct constraints and may enable morphological conservation amidst adaptation. Dev Biol. 2017. doi: 10.1016/j.ydbio.2017.07.019. PubMed PMID: 28780048.

18. Kalinka AT, Varga KM, Gerrard DT, Preibisch S, Corcoran DL, Jarrells J, et al. Gene expression divergence recapitulates the developmental hourglass model. Nature. 2010;468(7325):811–4. doi: 10.1038/nature09634. PubMed PMID: 21150996.

19. Piasecka B, Lichocki P, Moretti S, Bergmann S, Robinson-Rechavi M. The hourglass and the early conservation models--co-existing patterns of developmental constraints in vertebrates. PLoS Genet. 2013;9(4):e1003476. doi: 10.1371/journal.pgen.1003476. PubMed PMID: 23637639; PubMed Central PMCID: PMC3636041.

20. Levin M, Anavy L, Cole AG, Winter E, Mostov N, Khair S, et al. The mid-developmental transition and the evolution of animal body plans. Nature. 2016;531(7596):637–41. doi: 10.1038/nature16994. PubMed PMID: 26886793; PubMed Central PMCID: PMCPMC4817236.

21. Tu Q, Cameron RA, Davidson EH. Quantitative developmental transcriptomes of the sea urchin Strongylocentrotus purpuratus. Dev Biol. 2014;385(2):160–7. Epub 2013/12/03. doi: 10.1016/j.ydbio.2013.11.019. PubMed PMID: 24291147; PubMed Central PMCID: PMC3898891.

22. Tu Q, Cameron RA, Worley KC, Gibbs RA, Davidson EH. Gene structure in the sea urchin Strongylocentrotus purpuratus based on transcriptome analysis. Genome Res. 2012;22(10):2079–87. doi: 10.1101/gr.139170.112. PubMed PMID: 22709795; PubMed Central PMCID: PMCPMC3460201.

23. Materna SC, Nam J, Davidson EH. High accuracy, high-resolution prevalence measurement for the majority of locally expressed regulatory genes in early sea urchin development. Gene Expr Patterns. 2010;10(4-5):177–84. Epub 2010/04/20. doi: 10.1016/j.gep.2010.04.002. PubMed PMID: 20398801; PubMed Central PMCID: PMC2902461.

24. Young MD, Wakefield MJ, Smyth GK, Oshlack A. Gene ontology analysis for RNA-seq: accounting for selection bias. Genome Biol. 2010;11(2):R14. Epub 2010/02/06. doi: 10.1186/gb-2010-11-2-r14. PubMed PMID: 20132535; PubMed Central PMCID: PMC2872874.

25. Tsvia Gildor GC, Maya Lalzar, Veronica Hinman and Smadar Ben-Tabou de-Leon. Echinobase viewer of P. miniata quantitative developmental transcriptomes www.echinobase.org/shiny/quantdevPm.

26. Chang W, Cheng J, Allaire J, Xie Y, McPherson J. Framework for R. R package version 1.2.0. https://CRANR-projectorg/package=shiny 2018.

27. Cameron RA, Samanta M, Yuan A, He D, Davidson E. SpBase: the sea urchin genome database and web site. Nucleic Acids Res. 2009;37(Database issue):D750–4. Epub 2008/11/18. doi: gkn887 [pii] 10.1093/nar/gkn887. PubMed PMID: 19010966; PubMed Central PMCID: PMC2686435.

28. Barsi JC, Tu Q, Calestani C, Davidson EH. Genome-wide assessment of differential effector gene use in embryogenesis. Development. 2015;142(22):3892–901. doi: 10.1242/dev.127746. PubMed PMID: 26417044; PubMed Central PMCID: PMCPMC4712884.

29. Solek CM, Oliveri P, Loza-Coll M, Schrankel CS, Ho EC, Wang G, et al. An ancient role for Gata-1/2/3 and Scl transcription factor homologs in the development of immunocytes. Dev Biol. 2013;382(1):280–92. Epub 2013/06/25. doi: 10.1016/j.ydbio.2013.06.019. PubMed PMID: 23792116.

30. Flores RL, Livingston BT. The skeletal proteome of the sea star Patiria miniata and evolution of biomineralization in echinoderms. BMC Evol Biol. 2017;17(1):125. doi: 10.1186/s12862-017-0978-z. PubMed PMID: 28583083; PubMed Central PMCID: PMCPMC5460417.

31. She X, Rohl CA, Castle JC, Kulkarni AV, Johnson JM, Chen R. Definition, conservation and epigenetics of housekeeping and tissue-enriched genes. BMC Genomics. 2009;10:269. doi: 10.1186/1471-2164-10-269. PubMed PMID: 19534766; PubMed Central PMCID: PMCPMC2706266.

32. Zabidi MA, Arnold CD, Schernhuber K, Pagani M, Rath M, Frank O, et al. Enhancer-core-promoter specificity separates developmental and housekeeping gene regulation. Nature. 2015;518(7540):556–9. doi: 10.1038/nature13994. PubMed PMID: 25517091.

33. Bogdanovic O, Smits AH, de la Calle Mustienes E, Tena JJ, Ford E, Williams R, et al. Active DNA demethylation at enhancers during the vertebrate phylotypic period. Nat Genet. 2016;48(4):417–26. doi: 10.1038/ng.3522. PubMed PMID: 26928226; PubMed Central PMCID: PMCPMC5912259.

34. Gildor T, Ben-Tabou de-Leon S. Comparative Study of Regulatory Circuits in Two Sea Urchin Species Reveals Tight Control of Timing and High Conservation of Expression Dynamics. PLoS Genet. 2015;11(7):e1005435. doi: 10.1371/journal.pgen.1005435. PubMed PMID: 26230518.

35. Bolger AM, Lohse M, Usadel B. Trimmomatic: a flexible trimmer for Illumina sequence data. Bioinformatics. 2014;30(15):2114–20. doi: 10.1093/bioinformatics/btu170. PubMed PMID: 24695404; PubMed Central PMCID: PMCPMC4103590.

36. Haas BJ, Papanicolaou A, Yassour M, Grabherr M, Blood PD, Bowden J, et al. De novo transcript sequence reconstruction from RNA-seq using the Trinity platform for reference generation and analysis. Nat Protoc. 2013;8(8):1494–512. Epub 2013/07/13. doi: 10.1038/nprot.2013.084. PubMed PMID: 23845962.

37. Grabherr MG, Haas BJ, Yassour M, Levin JZ, Thompson DA, Amit I, et al. Full-length transcriptome assembly from RNA-Seq data without a reference genome. Nat Biotechnol. 2011;29(7):644–52. Epub 2011/05/17. doi: 10.1038/nbt.1883. PubMed PMID: 21572440; PubMed Central PMCID: PMC3571712.

38. Waterhouse RM, Seppey M, Simao FA, Manni M, Ioannidis P, Klioutchnikov G, et al. BUSCO applications from quality assessments to gene prediction and phylogenomics. Mol Biol Evol. 2017. doi: 10.1093/molbev/msx319. PubMed PMID: 29220515; PubMed Central PMCID: PMCPMC5850278.

39. Aubry S, Kelly S, Kumpers BM, Smith-Unna RD, Hibberd JM. Deep evolutionary comparison of gene expression identifies parallel recruitment of trans-factors in two independent origins of C4 photosynthesis. PLoS Genet. 2014;10(6):e1004365. doi: 10.1371/journal.pgen.1004365. PubMed PMID: 24901697; PubMed Central PMCID: PMCPMC4046924.

40. Bray NL, Pimentel H, Melsted P, Pachter L. Near-optimal probabilistic RNA-seq quantification. Nat Biotechnol. 2016;34(5):525–7. doi: 10.1038/nbt.3519. PubMed PMID: 27043002.

41. Soneson C, Love MI, Robinson MD. Differential analyses for RNA-seq: transcript-level estimates improve gene-level inferences. F1000Res. 2015;4:1521. doi: 10.12688/f1000research.7563.2. PubMed PMID: 26925227; PubMed Central PMCID: PMCPMC4712774.

42. Love MI, Huber W, Anders S. Moderated estimation of fold change and dispersion for RNA-seq data with DESeq2. Genome Biol. 2014;15(12):550. doi: 10.1186/s13059-014-0550-8. PubMed PMID: 25516281; PubMed Central PMCID: PMCPMC4302049.

43. Dobin A, Davis CA, Schlesinger F, Drenkow J, Zaleski C, Jha S, et al. STAR: ultrafast universal RNA-seq aligner. Bioinformatics. 2013;29(1):15–21. doi: 10.1093/bioinformatics/bts635. PubMed PMID: 23104886; PubMed Central PMCID: PMCPMC3530905.

44. Anders S, Pyl PT, Huber W. HTSeq--a Python framework to work with high-throughput sequencing data. Bioinformatics. 2015;31(2):166–9. doi: 10.1093/bioinformatics/btu638. PubMed PMID: 25260700; PubMed Central PMCID: PMCPMC4287950.

45. Robinson MD, McCarthy DJ, Smyth GK. edgeR: a Bioconductor package for differential expression analysis of digital gene expression data. Bioinformatics. 2010;26(1):139–40. Epub 2009/11/17. doi: 10.1093/bioinformatics/btp616. PubMed PMID: 19910308; PubMed Central PMCID: PMC2796818.

