## Supplemental figures for "Developmental transcriptomes of the sea star, *Patiria miniata*, illuminate the relationship between conservation of gene expression and morphological conservation"

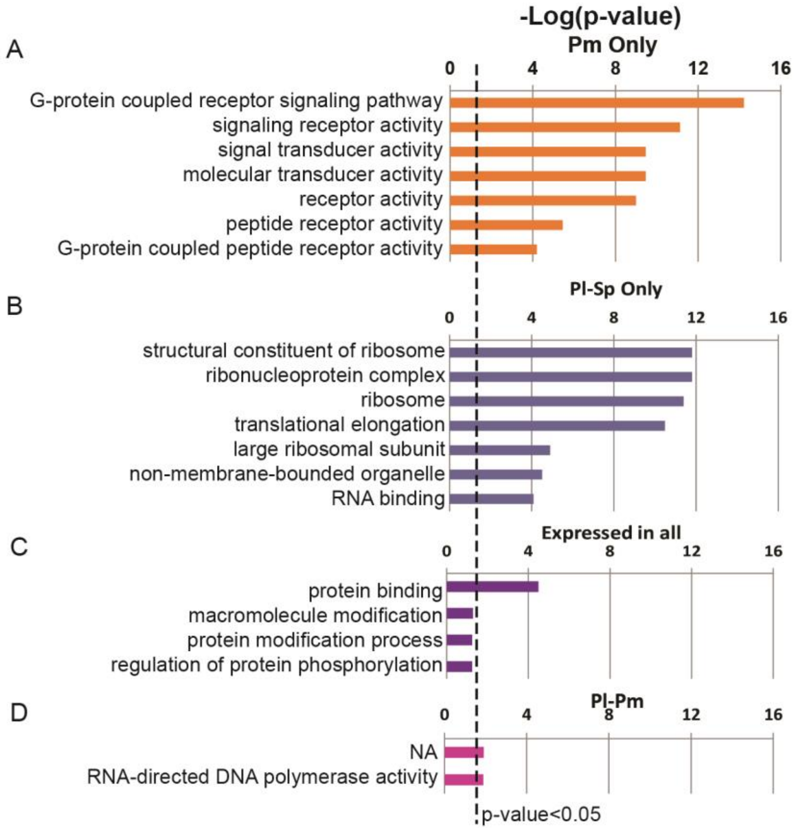


**Figure S1 GO enrichment in the different expression groups.** A, GO enrichment in genes expressed only in *P. miniata*, B, GO enrichment in gene expressed only in the two sea urchins, C, GO enrichment in genes that are expressed in the three species, D, GO enrichment in genes expressed in only in *P. lividus* and *P. miniata*.

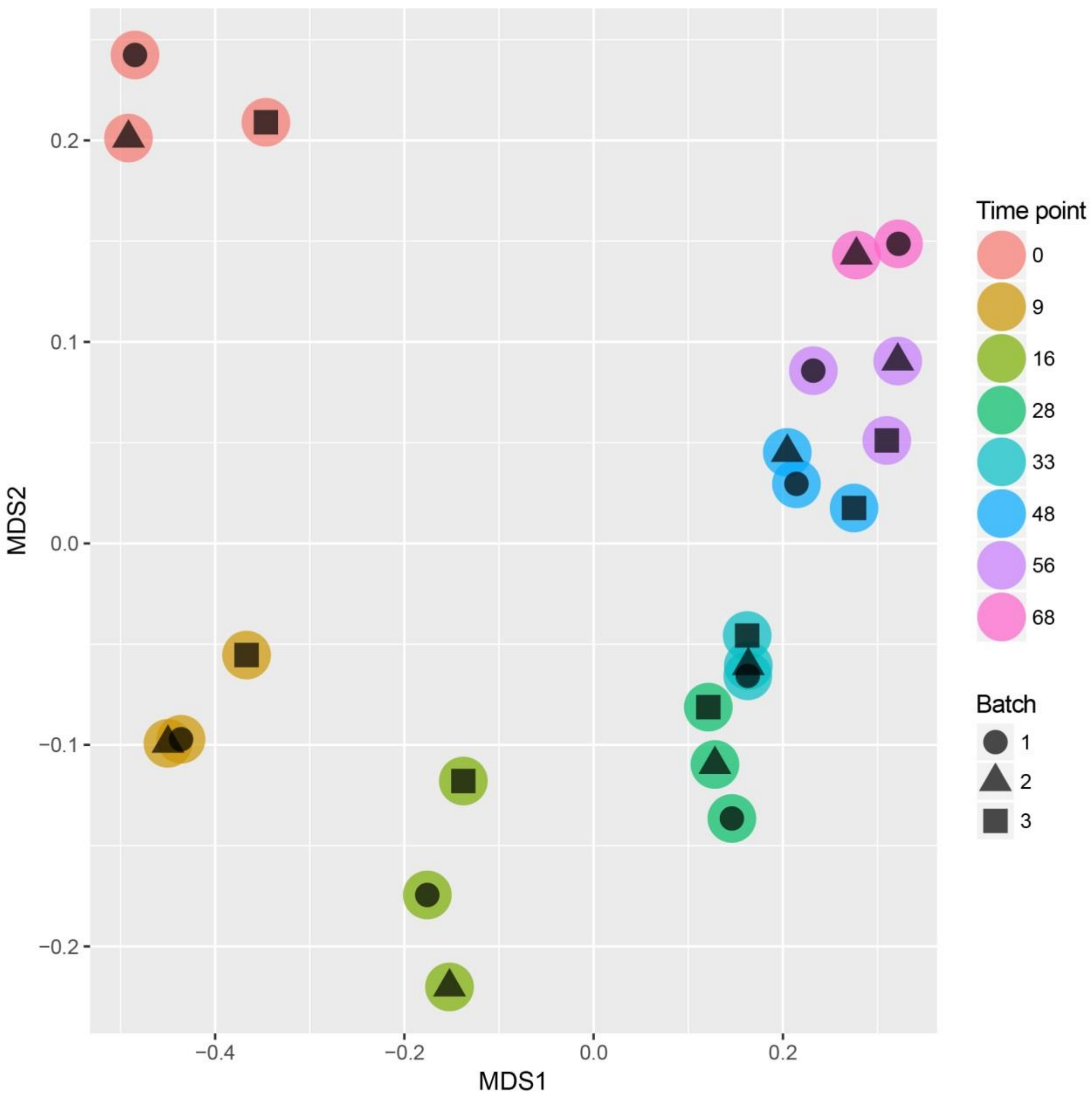


**Figure S2 NMDS analysis of the sea star expression profiles for the three biological replicates show high reproducibility of gene expression.** NMDS of individual *P. miniata* time points and biological replicates. The different batches are indicated as circles, triangles and squares and different developmental time points are indicated by color.

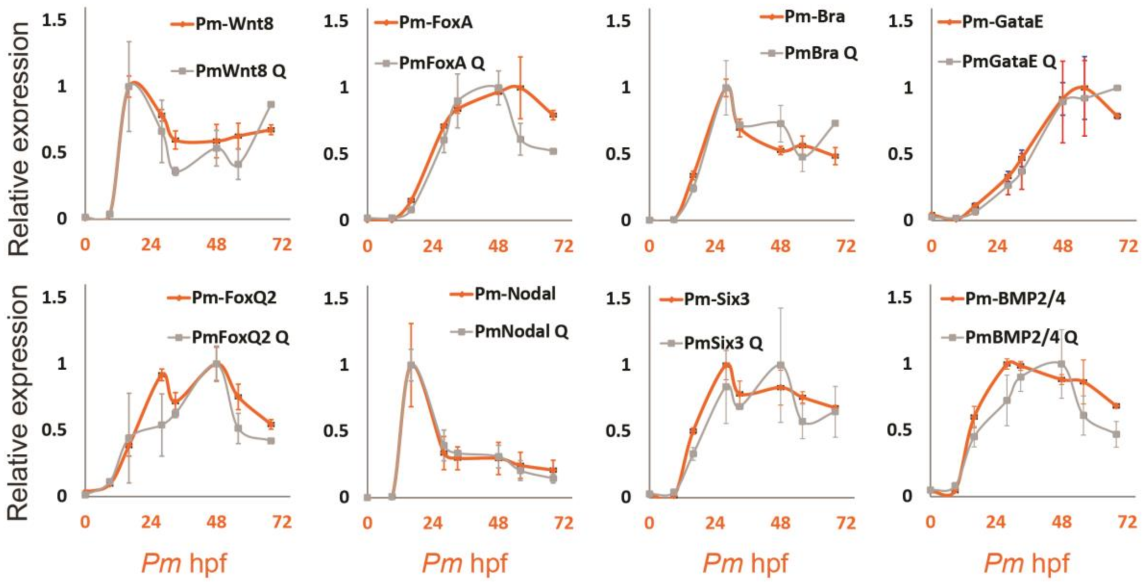


**Figure S3 QPCR verification of RNA-seq results shows high agreement between the two measurements.** Orange curves in each graph show RNA-seq results and grey lines show QPCR measurements at the same time points taken from [1]. Error bars correspond to standard deviation of three biological replicates in both experiments. Gene's name is indicated in each panel.

1. Gildor T, Hinman V, Ben-Tabou-De-Leon S. Regulatory heterochronies and loose temporal scaling between sea star and sea urchin regulatory circuits. Int J Dev Biol. 2017;61(3-4-5):347-56. doi: 10.1387/ijdb.160331sb. PubMed PMID: 28621432.
